# Common garden experiment reveals altered nutritional values and DNA methylation profiles in micropropagated three elite Ghanaian sweet potato genotypes

**DOI:** 10.1101/471623

**Authors:** Belinda Akomeah, Marian D. Quain, Sunita A. Ramesh, Carlos M. Rodríguez López

**Affiliations:** ARC Centre of Excellence in Plant Energy Biology, University of Adelaide, Waite Campus, PMB1 Glen Osmond, SA, 5064; The Waite Research Institute and The School of Agriculture, Food and Wine, University of Adelaide, Waite Campus, PMB1 Glen Osmond, SA, 5064; CSIR-Crops Research Institute, P. O. Box 3785, Kumasi, Ghana; Environmental Epigenetics and Genetics Group, Department of Horticulture, College of Agriculture, Food and Environment, University of Kentucky, Lexington, KY, USA

## Abstract

Micronutrient deficiency is the cause of multiple diseases in developing countries. Staple crop biofortification is an efficient means to combat such deficiencies in the diets of local consumers. Biofortified lines of sweet potato (*Ipomoea batata* L. Lam) with enhanced beta-carotene content have been developed in Ghana to alleviate Vitamin A Deficiency. These genotypes are propagated using meristem micropropagation to ensure the generation of virus-free propagules. *In vitro* culture exposes micropropagated plants to conditions that can lead to the accumulation of somaclonal variation with the potential to generate unwanted aberrant phenotypes. However, the effect of micropropagation induced somaclonal variation on the production of key nutrients by field-grown plants has not been previously studied. Here we assessed the extent of *in vitro* culture induced somaclonal variation, at a phenotypic, compositional and genetic/epigenetic level, by comparing field-maintained and micropropagated lines of three elite Ghanaian sweet potato genotypes grown in a common garden. Although micropropagated plants presented no observable morphological abnormalities compared to field maintained lines, they presented significantly lower levels of iron, total protein, zinc, and glucose. Methylation Sensitive Amplification Polymorphism analysis showed a high level of *in vitro* culture induced molecular variation in micropropagated plants. Epigenetic, rather than genetic variation, accounts for most of the observed molecular variability. Taken collectively, our results highlight the importance of ensuring the clonal fidelity of the micropropagated biofortified lines in order to reduce potential losses in the nutritional value prior to their commercial release.

## Introduction

Sweet potato (*Ipomoea batatas* L. Lam), is a drought tolerant, low input, and high yielding crop, which produces more nutrients and has higher edible energy than most staples such as rice, cassava, wheat, and sorghum [1]. As a predominantly vegetatively propagated crop, virus accumulation in vegetative propagules (i.e. vine cuttings and tubers) can cause devastating loss in yield and poor root quality in subsequent cultivation [2]. Micropropagation techniques, such as meristem or nodal tip culture, coupled with thermotherapy or cryotherapy, are currently the principal plant tissue culture (PTC) methods for producing healthy (pathogen-tested/disease-free) clones of planting materials [3]. However, the generation of true-to-type material through *in vitro* propagation can be challenging due to somaclonal variation [4].

Somaclonal variation refers to changes that can be induced during *in vitro* tissue culture and have been reported in all *in vitro* systems [5-8]. Such changes can be genetic and/or epigenetic in nature. Epigenetic modifications are heritable changes that can affect the phenotype without changes to the DNA sequence [9]. These are mediated, among other mechanisms, by DNA methylation, small RNA mediated silencing, histone modification, and chromatin remodelling [10]. DNA methylation is the addition of a methyl group to carbon 5 in the pyrimidine ring of cytosines [11]. In plants, DNA methylation occurs at the CG, CNN, or CNG context (where N=C, A, or T), and has been shown to induce changes in gene expression, which has the potential to lead to phenotypic changes [12]. Environmental conditions can induce changes to plant methylomes [13, 14]. *In vitro* culture of plant tissues has been reported to induced epigenetic somaclonal variation for multiple crop species including garlic [15], cassava [6], pineapple [16], cotton [17], cocoa [7], and other crops [5, 18]. However, few studies have evaluated the extent of DNA methylation changes during meristem propagation of sweetpotato. In addition, no study has been conducted to understand the correlation between the extent of *in vitro* induced epigenetic changes and the nutritional composition of sweet potato tubers.

Vitamin A is an essential nutrient that is required in small amounts for maintaining healthy growth and development, particularly in growing children, pregnant and lactating mothers [19]. Vitamin A deficiency (VAD) has been declared a public health problem affecting up to 48% of children in sub-Saharan African countries including Ghana [20]. VAD manifests itself as severe respiratory infections, diarrhoeal diseases and eye diseases ranging from night blindness, to the more serious sight condition, keratomalacia (melting of the cornea) and even mortality [21]. Beta carotene is a precursor to Vitamin A abundant in plant cells [22]. Biofortification is an affordable tool to combat nutrient deficiencies and hence sweet potato is currently being biofortified for enhanced beta carotene content to combat micronutrient malnutrition. It is, however, crucial to understand the impact that somaclonal variation via micropropagation has on the nutritional content of biofortified sweet potato tubers. Thus, our aim was to test the hypothesis that the nutritional value of micropropagated plants could be affected by somalconal variation. To achieve this we assesed differences in plant morphology and nutritional composition of *in vitro* and field-maintained propagules of three improved sweet potato genotypes, while evaluating the incidence of molecular (genetic and epigenetic) somaclonal variation. Micropropagated and field maintained plants of the selected sweet potato genotypes were grown in a common garden experiment and examined for phenotypic variation in the micropropagated regenerants. Near Infrared Spectrophotometry (NIRS) [23] was used to analyse the nutritional composition in mature tubers from both types of propagules. Finally, Methylation Sensitive Amplification Polymorphism (MSAP), which is a rapid, cost effective, and reliable method of assessing epigenetic variability [24], was used to investigate the extent of genetic and epigenetic variability imposed by *in vitro* culture on micropropagated plants.

## Materials and methods

### Field experimental design

Field work for this study was conducted during the major growing season (March-July, 2016), at the Council for Scientific and Industrial Research-Crops Research Institute (CSIR-CRI), Fumesua, located in the Forest agro-ecological zone of Ghana (N 6.43’25°, W 1.31’9°). The land was cleared, ploughed, ridged, and manured with poultry droppings [25]. Three CRI improved genotypes with moderate resistance to sweet potato viruses were used for this study: CRI-Bohye, CRI-Ogyefo, and CRI-Otoo (Table S1). Micropropagated clones of these genotypes, produced by meristem-tip culture and thermotherapy and maintained for 18 months in Plant Tissue Culture [3, 26], were obtained from the screen house. Cuttings from visually virus free planting vines of the same genotypes maintained according to the agronomic practices of the institute on the CSIR-CRI multiplication field (field-maintained), since year of release (Table S1), were also obtained. Both types of propagules were planted in a Randomized Complete Block Design with three replicated blocks (Fig. S1). Eight plants per plot were randomly selected for molecular, morphological and nutritional analysis.

### Phenotypic characterisation

To examine the incidence of phenotypic somaclonal variation, all plants were scored on a scale of 1-9 for selected foliage and storage roots characteristics based on standard sweet potato morphological descriptors [27] (Table S2). Selected descriptors were based on pigmentation in the leaves, roots, and vines. The foliage parameters were scored between 90-100 days after planting, and they included immature and mature leaf colour, abaxial leaf vein pigmentation, predominant vine colour, secondary vine colour, petiole pigmentation, and plant type. Storage root descriptors, i.e. shape, predominant skin colour, and flesh colour were documented at 120 days after planting [27].

### Nutritional analysis

To study the tissue culture induced changes to the composition of mature tubers, nutritient content analysis of the storage roots was done at harvest. Analysis was carried out at the Quality and Nutritional laboratory, CSIR-CRI. In brief, harvested roots were pooled by block/genotype/propagation system (n=3). Each pool was then sampled individually as described by Amankwaah [28]. Tubers were then washed, air dried, peeled, quartered, sliced, weighed (50g), and freeze dried for 73 hours using a YK-118 Vacuum Freeze Dryer (True Ten Industrial Company Limited Taichung, Taiwan). Freeze-dried weights were recorded, and dry matter was computed based on the differences between the fresh and freeze-dried weights as: Percentage dry matter = dry weight/fresh weight × 100. Freeze-dried tuber samples were then milled (3383-L70, Thomas Scientific, Dayton Electric Manufacturing Company Limited, IL 60714, USA), and analysed using Near Infrared Spectrophotometry (NIRS) (XDS Rapid Content Analyzer, Hoganae, Sweden) to estimate starch (%), protein (%), zinc (mg 100g^-1^), fructose (%), glucose (%), iron (mg 100g^-1^) and sucrose (%) content [29].

## Analysis of plant DNA methylation profiles

### DNA extraction

In all, 144 plants were sampled for DNA extraction, comprising of 48 samples for each of the three genotypes (24 micropropagated, 24 field-maintained). The youngest leaves of 6 weeks old plants were collected on ice from the field, and kept in liquid nitrogen until DNA extraction with the modified CTAB protocol [30]. Agarose gel electrophoresis (0.8%) was used to check the quality of the extracted DNA, while the concentration and purity were analysed using NanoDrop 1000 Spectrophotometer (Thermo Scientific, Wilmington, USA). The DNA was diluted to 20 ng μl^-1^ for MSAP analysis.

### Methylation Sensitive Amplification Polymorphism (MSAP) Profiling

To investigate the tissue culture induced changes to cytosine methylation, MSAP was performed based on established protocol [31]. Genomic DNA was digested with a combination of one of two methylation sensitive isoschizomers as frequent cutter (i.e. *Hpa*II or *Msp*I which present the same recognition sequence (CCGG), but different sensitivity to DNA methylation), and the methylation insensitive *Eco*RI as rare cutter, which has the recognition site GAATTC. Restriction products were then ligated to double stranded DNA adaptors with co-adhesive ends complementary to those present in *Hpa*II/*Msp*I and *Ec*oRI restriction products using T4 DNA ligase. Pre-selective amplification was then done using primers complimentary to the adaptor sequence, but with unique 3’overhangs (Table S3). A second round of selective amplification was then carried out using primers with extra selective bases and labelled with a 6-FAM reporter molecule for fragment detection (Table S3). Finally, amplified products were capillary electrophoresed using the ABI PRISM^®^ 3130 Genetic Analyzer (Applied Biosystems, Foster City, CA, USA) at the Australian Genome Research Facility Ltd, Adelaide, South Australia. MSAP capillary electrophoresis profiles were transformed into a presence (1) or absence (0) binary matrix for both *Hpa*II/*Eco*R1 and *Msp*I/*Eco*R1 restriction products, using GeneMapper Software v4 (Applied Biosystems, Foster City, CA).

To identify the most informative MSAP primer combinations for this study, twelve selective primer combinations (Table S4) were tested on three micropropagated and three field-maintained DNA samples from one genotype (Bohye). One of the three samples of each group was duplicated to assess marker reproducibility for each primer combination. Primer reproducibility (i.e. % of MSAP markers present in both replicates), number of alleles, number of differential alleles (i.e. number of polymorphic markers between field maintained and micropropagated samples), and principal coordinate analysis (PCoA) of MSAP profiles of all the primer combinations were analysed to determine the two most informative primer pairs that were then used in all samples.

### Statistical analysis

To examine phenotypic, nutrient content, and virus incidence data, GenStat (15^th^ Edition) was used to perform Analysis of Variance (ANOVA), and to statistically test any detected difference between means at 5% least significant difference (LSD).

GenAlEx v6.5 [32] and R package *msap* v.3.3.1 [24] were used to investigate the level of tissue culture induced genetic and epigenetic variation by analysing both types of variability among and between field-maintained and micropropagated samples for each of the 3 genotypes tested as described previously [13, 31]. In brief, first we used Principal Coordinate Analysis (PCoA) in GenAlEx v6.5 to visualise the molecular diversity (genetic and epigenetic) captured by MSAP profiles generated with each primer (E and I) and enzyme combination (*Hpa*II/*EcoR*I and *Msp*I *EcoR*I) individually. Then Analysis of Molecular Variance (AMOVA) was computed on GenAlEx v6.5 to estimate pairwise molecular distances (PhiPT) between the field-maintained and micropropagated plant populations. The significance of the observed PhiPT values was assigned by random permutations tests (based on 9,999 replicates).

To determine to what extent the observed changes in MSAP profiles could be attributed to genetic (changes in DNA sequence) or epigenetic (changes in the DNA methylation patterns) of the studied samples, we identified Non-Methylated Loci (NML) and Methylation Susceptible Loci (MSL) by comparing the MSAP profiles generated using *Hpa*II/*EcoR*I and *Msp*I *EcoR*I as implemented in *msap v.3.3.1*. First, Principal Component Analysis (PCA) was used to visualise the contribution of each type of change, then Shannon diversity Index (S) and Wilcoxon Rank Sum test were used to estimate the contribution and statistical significance of each type of variability (genetic and epigenetic).

## Results

### Morphological characterisation

No significant variation was observed for any of the 8 phenotypic traits measured i.e. foliage colour (for immature and mature plants), plant type, petiole pigmentation, abaxial leaf vein pigmentation, storage root shape, storage root pigmentation, and flesh colour between micropropagated and field-maintained plants in any of the genotypes analysed (Table S5).

### Nutritional composition

Analysis of variance of eight nutrients in mature sweet potato tubers showed that iron, total protein, zinc, and glucose levels were significantly higher (P<0.05) in field-maintained than in micropropagated sweet potato plants (Fig 1, Table 1). Sucrose, fructose, dry matter, and starch contents were not significantly different between micropropagated and field propagules for the genotypes analysed (Table 1). For all measured traits (excluding sucrose and zinc), variability was higher in micropropagated samples than in field maintained ones (Fig 1, Table 1).

**Table 1.** Compositional analysis of mature tubers from field maintained (Field) and micropropagated (*in vitro*) sweet potato plants. Values show the mean values and standard deviation (in parenthesis) of nutrients in sweet potato tubers from three genotypes micropropagated and field maintained. Three replicate measurements were taken for each genotype/propagation system combination (n = 9). l.s.d.: least significant difference; n.s.: not significant; * P<0.05; ** P<0.005.

| Prop system | Dry Matter % | Fructose % | Glucose % | Iron mg/100g | Protein % | Starch % | Sucrose % | Zinc mg/100g |
| --- | --- | --- | --- | --- | --- | --- | --- | --- |
| Field | 30.64<br>(2.37) | 1.32<br>(0.31) | 2.74<br>(0.31) | 2.46<br>(0.16) | 5.95<br>(1.02) | 67.45<br>(1.16) | 5.61<br>(4.60) | 1.31<br>(0.21) |
| *In vitro* | 31.96<br>(3.46) | 1.38<br>(0.47) | 2.36<br>(0.55) | 1.91<br>(0.37) | 4.68<br>(1.39) | 69.56<br>(4.21) | 4.71<br>(2.06) | 0.98<br>(0.20) |
| l.s.d (5%) | n.s. | n.s. | 0.17** | 0.26** | 1.10* | n. s | n. s | 0.20* |

**Fig 1.**
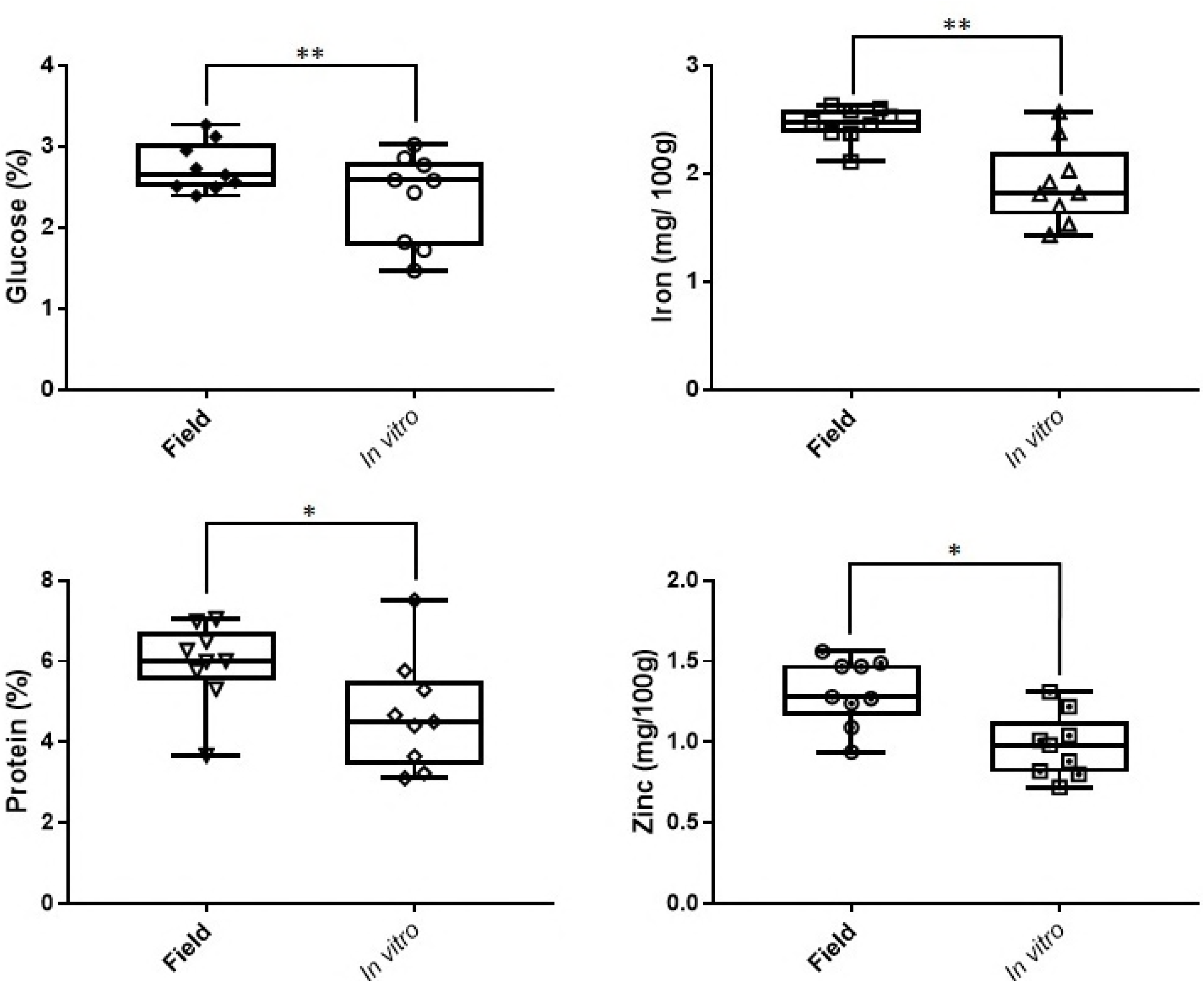
Effect of somaclonal variation on nutritional composition of sweet potato tubers. Box and whisker plot showing the content of glucose, iron, protein, and zinc in *in vitro* and field-maintained sweet potato tubers. Micropropagated and field maintained plants from three genotypes (Bohye, Ogyefo, and Otoo) were grown on a Randomized Complete Block Design with three replicated blocks and 24 plants per block/genotype/propagation system. Plants from each block/genotype/propagation system were pooled and analysed using Near Infrared Spectrophotometry.

### Assessment of micropropagation induced molecular variability

We first used a reduced number of samples to test 12 MSAP primer combinations to identify the most informative and reproducible primer combinations. These generated between 149 and 205 loci (Combinations K and D respectively) (Table S4). Estimated loci reproducibility ranged from 91 to 98% (Combinations G and L) (Table S4). The number of loci discriminating between field-maintained and micropropagated samples varied between 1 and 12 (Combinations H and C respectively) (Table S4). After comparison of the principal coordinates analysis results (Fig. S2), number of alleles, reproducibility, and number of discriminatory alleles between *in vitro* and field maintained plants for each of the 12 primer combinations, primer combinations E and I (Table S4) were selected to analyse differences between the entire sample set of *in vitro* and field maintained plants. These produced 197 and 174 alleles, 9 and 11 discriminatory alleles, and a reproducibility of 98% and 93% respectively.

When applied to all samples, primer combinations E and I generated a total of 244 and 235 loci respectively. PCoA and PCA of MSAP profiles containing all 479 loci showed that *in vitro* maintained samples, irrespective of their genotype, shift in the same direction, e.g. MSAP profiles generated from micropropagated plants, using both *Hpa*II and *Msp*I and irrespective of their background genotype, are displaced towards the top quadrants in relation to the field maintained plants, when analysed with GenAlex v6.5 (Fig. S3a) or towards the right quadrants when analysed using *msap* v.3.3.1 (Fig. S3b).

PCoA (as implemented on GenAlex v6.5) of MSAP profiles from samples grouped by genotype, showed clear separation between the populations of micropropagated and field maintained samples for each of the three genotypes studied (Fig 2). Analysis of the Molecular Variance (AMOVA) showed that differences between micropropagated and field maintained plants explain 13, 27 and 29% and 7, 22, and 24% of total variability observed for Ogyefo, Otoo, and Bohye for *Msp*I and *Hpa*II restriction products respectively. Pairwise molecular distances (PhiPT) calculated between micropropagated and field-maintained plant populations showed that all genotype/enzyme/primer combinations generated significant differences (P value < 0.005). Of these, micropropagated and field-maintained Ogyefo plants showed lower levels of molecular differentiation than those shown by Bohye and Otoo plants (Table 2). In all genotype/enzyme/primer combinations, micropropagated samples occupied a much larger Eigen space than their field-maintained counterparts (Fig 2).

**Fig 2.**
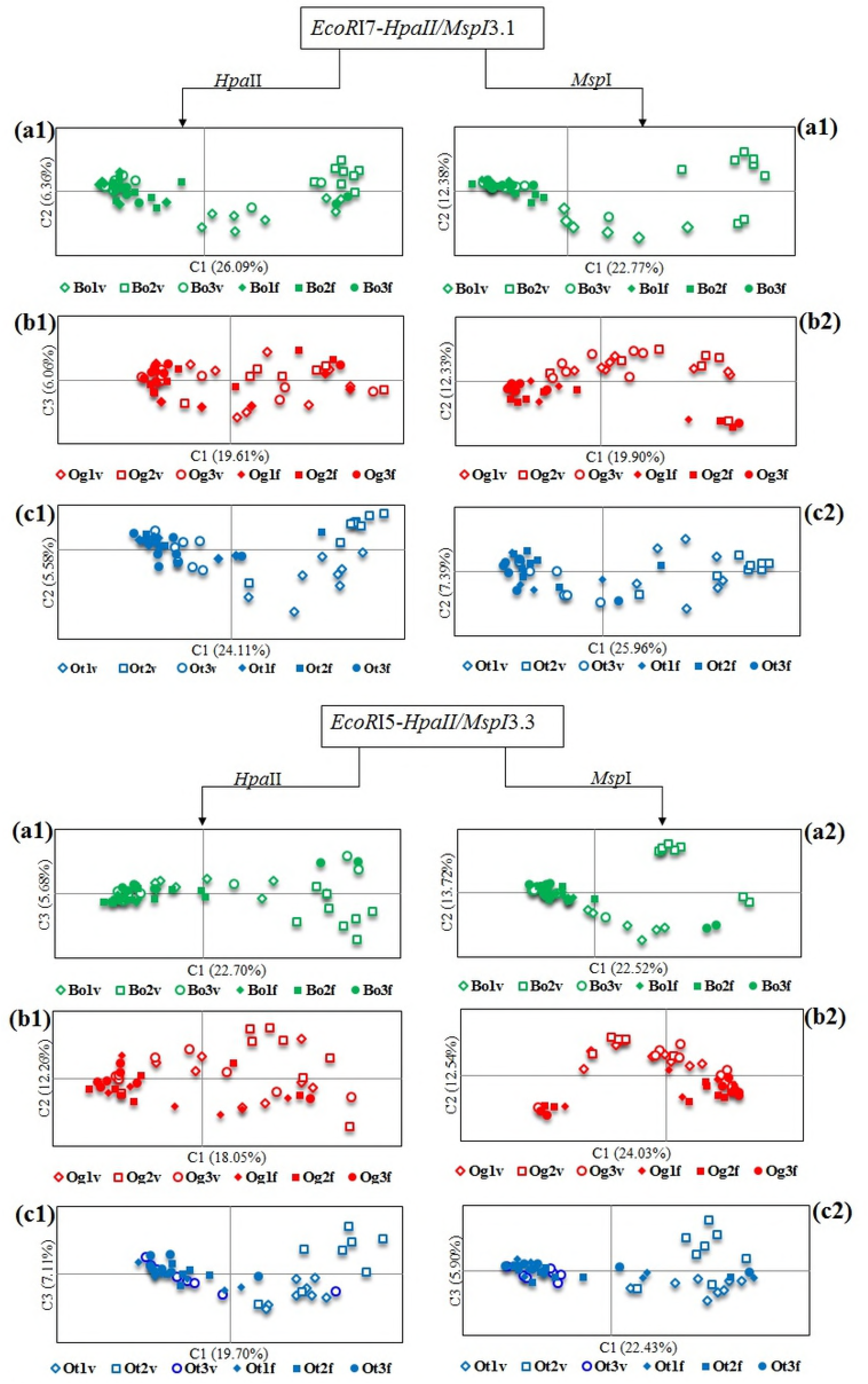
Analysis of somaclonal variation induced by micropropagation in three sweet potato genotypes. PCoA generated using GenAlex v6.5 from MSAP profiles from 144 micropropagated (empty symbols) and field-maintained (full symbols) plants from genotypes of Bohye (green), Ogyefo (red), and Otoo (blue) (n = 24). MSAP profiles were amplified from genomic DNA restricted using *Hpa*II (a1-f1) and *Msp*I (a2-f2) and amplified using primer combinations E (a-c) and I (d-f). Plants were grown on a Randomized Complete Block Design with three replicated blocks and 24 plants/block/genotype/propagation system.

**Table 2.** Analysis of molecular differentiation between field maintained and micropropagated sweet potato plants. Molecular distance (PhiPT) was calculated as implemented in GenAlex v6.5 using MSAP profiles generated from DNA from field maintained and micropropagated plants of three sweet potato genotypes (Bohye, Ogyefo, and Otoo) (n=24) restricted using *Hpa*II and *Msp*I and amplified using primer I and E. P values were calculated by random permutations tests based on 9,999 replicates.

|  | *Primer combination I* |  |  |  | *Primer combination E* |  |  |  |
| --- | --- | --- | --- | --- | --- | --- | --- | --- |
|  | *MspI* |  | *HpaII* |  | *HpaII* |  | *MspI* |  |
| Genotypes | PhiPT | P-value | PhiPT | P-value | PhiPT | P-value | PhiPT | P-value |
| Bohye F/M | 0.289 | 0.0001 | 0.218 | 0.001 | 0.168 | 0.0010 | 0.264 | 0.001 |
| Ogyefo F/M | 0.133 | 0.0001 | 0.068 | 0.002 | 0.088 | 0.0001 | 0.106 | 0.001 |
| Otoo F/M | 0.266 | 0.0001 | 0.241 | 0.001 | 0.166 | 0.0001 | 0.196 | 0.001 |

We then used *msap* v.3.3.1 R package to determine the contribution of epigenetic (Methylation-Susceptible Loci (MSL)) and of genetic (Non-Methylated Loci (NML)) to *in vitro* culture induced variability. PCA analysis of MSL and NML generated using both primer combinations showed separation between field-maintained and micropropagated plants (Fig S4). Pairwise PhiST distances between *in vitro* culture and field maintained plants showed epigenetic distances (i.e. PhiST calculated using MSL) were higher than those calculated using NML (i.e. genetic PhiST) for all genotypes (Table 3). As seen with PhiPT values, pairwise PhiST distances calculated for MSL and NML revealed that Ogyefo recorded the lowest epigenetic and genetic distances between micropropagated and field maintained plants, while Bohye presented the highest epigenetic distance, and Otoo had the highest genetic distance (Table 3). Shannon diversity Index (S) performed for MSL were higher (Paired sample T-test Pval<0.0001) than those for NML (Table 3). Wilcoxon Rank Sum test with continuity correction, which tests the significance of the Shannon Index, confirmed that the SIs calculated for MSL were statisticaly significant (P < 0.0001) while those calculated for NML were not (Table 3).

**Table 3.** Contribution of epigenetic and genetic polymorphisms to the molecular differentiation in sweet potato plants. MSL and NML were identified implementing msap package in R to MSAP profiles generated from DNA from field maintained and micropropagated plants of three sweet potato genotypes (Bohye, Ogyefo, and Otoo) (n=24) restricted using *Hpa*II and *Msp*I and amplified using primer E and I. #: number of loci; SI: Shannon Index; NA: not applicable; ^†^Wilcoxon rank sum test with continuity correction showing P < 0.0001; * P<0.0001.

| Primer combination E |  |  |  |  |  |  |  |  |
| --- | --- | --- | --- | --- | --- | --- | --- | --- |
|  | Methylation-Susceptible Loci (MSL) |  |  |  | No Methylated Loci (NML) |  |  |  |
|  | <sup>1</sup># | PhiST | P-value | SI | # | PhiST | P-val | SI |
| All genotypes | 181 | <sup>4</sup>NA | 0.0001* | 0.50469<sup>†</sup> | 63 | NA | 0.0001* | 0.23183 |
| Bohye | 172 | 0.113 | 0.0001* | 0.47136<sup>†</sup> | 72 | 0.107 | 0.0001* | 0.20482 |
| Ogyefo | 152 | 0.085 | 0.0001* | 0.50486<sup>†</sup> | 92 | 0.027 | 0.0595 | 0.25226 |
| Otoo | 151 | 0.097 | 0.0001* | 0.52322<sup>†</sup> | 93 | 0.115 | 0.0001* | 0.25464 |
| Primer combination I |  |  |  |  |  |  |  |  |
|  | Methylation-Susceptible Loci |  |  |  | No Methylated Loci |  |  |  |
|  | # | PhiST | P-val | SI | # | PhiST | P-val | SI |
| All genotypes | 171 | NA | 0.0001* | 0.5082<sup>†</sup> | 64 | NA | 0.0001* | 0.2098 |
| Bohye F/M | 153 | 0.235 | 0.0001* | 0.4939<sup>†</sup> | 82 | 0.052 | 0.0009* | 0.2048 |
| Ogyefo F/M | 145 | 0.130 | 0.0001* | 0.4998<sup>†</sup> | 90 | 0.039 | 0.0029* | 0.2333 |
| Otoo F/M | 152 | 0.135 | 0.0001* | 0.5131<sup>†</sup> | 83 | 0.179 | 0.0001* | 0.2290 |

Analysis of differences in band presence/absence between samples restricted with *Hpa*II or *Msp*I was used to infer the contribution of each type of methylation present on the enzymes recognition site (i.e CCGG). When considering all samples collectively (Fig 3), fully methylated recognition sites (i.e. all cytosines methylated) represented the majority of the analysed loci (48.2%), followed by fully unmethylated sites (21.5%), hemimethylated sites (i.e. only one DNA strand methylated presenting cytosines) (19.3%), and sites presenting internal cytosine methylation (11.2%). DNA from micropropagated plants presented lower levels of unmethylated (19.1 vs 23.9%) and hemimethylated sites (18.1 vs 20.5%). Micropropagated plants also showed higher levels of fully methylated sites (49.4 vs 47.1%) and of internal cytosine methylation (13.4 vs 9.04%) (Fig 3). Individual analysis by genotype revealed that Ogyefo propagules presented lowest levels on differentiation between micropropagated and field maintained plants (Fig. S4).

**Figure 3.**
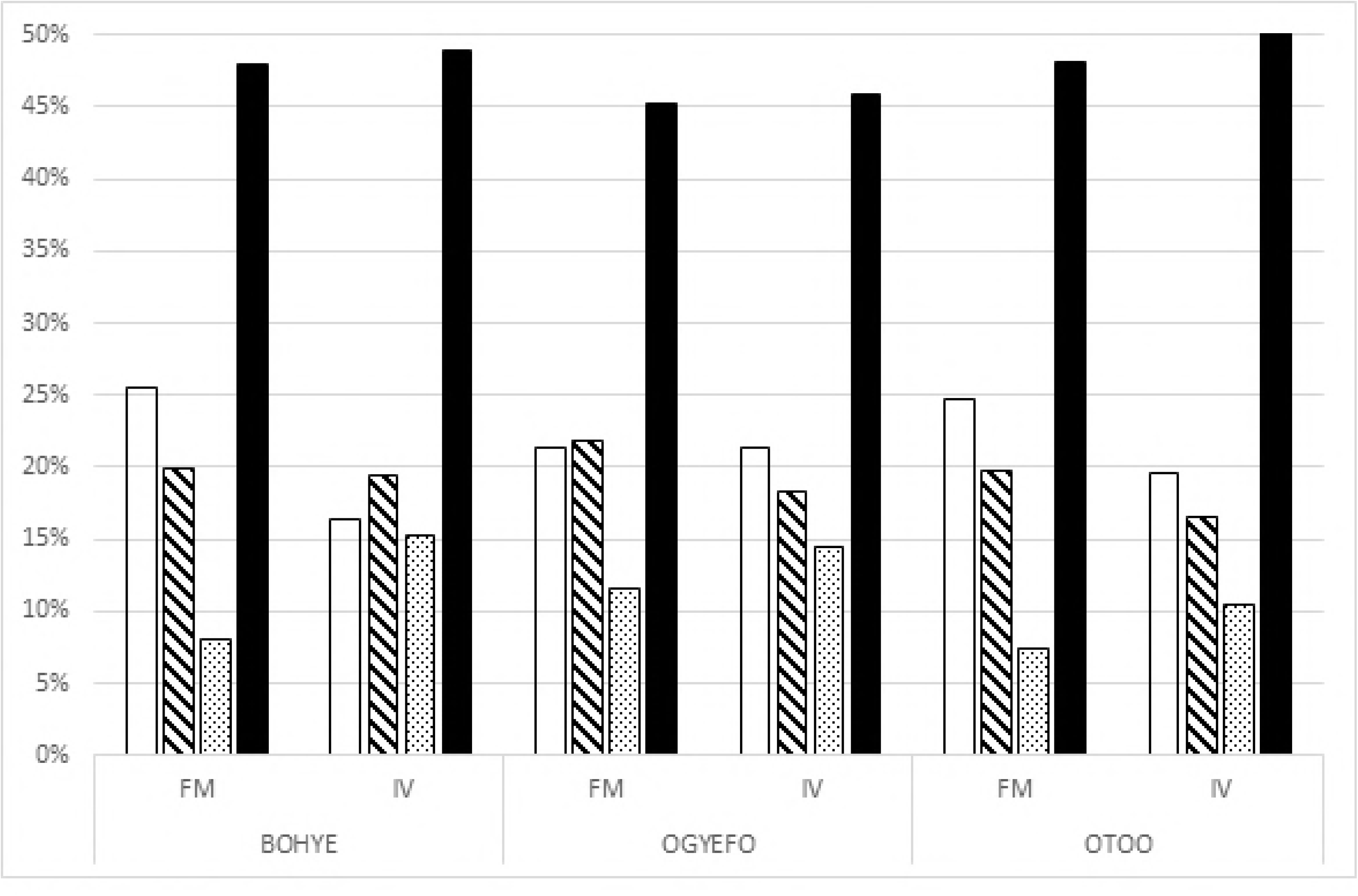
Analysis of somaclonal variation in sweet potato. Percentage of unmethylated loci (empty bars), hemimethylated loci (bars with diagonal pattern), loci containing internal methylation (dotted bars) and fully methylated loci (black bars) on MSAP profiles generated from micropropagated (VF) and field samples (FM) (n=24) from three sweet potato genotypes (Bohye, Ogyefo, Otoo) as determined by *msap* package in R.

## Discussion

The passage of plant tissues through *in vitro* culture may induce undesired variability in the regenerated propagules called somaclonal variation [10]. However, in some cases, *in vitro* culture is indispensable for the production and multiplication of disease-free planting materials in vegetatively propagated crops like sweet potato [33]. To test to what extent *in vitro* culture induced somaclonal variation could affect the nutritional value of micropropagated sweet potatoes, we compared field maintained morphological, chemical (nutritional composition) and molecular (genetic and epigenetic) variation in micropropagated sweet potato mericlones.

### Micropropagation alters sweet potato nutritional value

Phenotypic characterization, based on standard sweet potato morphological descriptors of micropropagated regenerants and their field-maintained counterparts, grown on a common garden setup, did not show any significant differences between both types of propagules. Conversely, compositional comparative analysis of mature tubers showed that 4 out of the 8 nutritional traits analysed (i.e. iron, zinc, total protein, and glucose) were significantly lower in micropropagated plants compared to field-maintained individuals. Also, the variability between samples in 6 out of the 8 measured nutrients (dry matter, fructose, glucose, iron, total protein, and starch), was higher in micropropagated plants than in field maintained plants. Previous studies have shown that the components of the growth media that are supplemented to tissue culture plants can have an effect on *in vitro* plants [6]. In this study, however, both micropropagated and field-maintained plants were grown in the same conditions from planting until harvest (four months). This indicates that the detected differences in tuber composition could be associated to somaclonal variability induced during culture that is maintained after plant establishment in the field. Both genetic and epigenetic somaclonal variability can be faithfully maintained during multiple cell divisions [6, 7], and therefore could be the source of the tuber compositional differences observed here.

### Micropropagation induces somaclonal variation in sweet potato

To determine the level and nature of somaclonal variation induced by micropropagation we analysed the MSAP profiles of 144 plants (i.e. 48 samples for each of the three genotypes (24 micropropagated, 24 field-maintained)), grown in a common garden set up. Multivariate analysis (PCoA and PCA) revealed that MSAP profiles of micropropagated plants are different from those of generated from field maintained plants. Analysis of Molecular Variance (AMOVA) showed that between 7 and 29% of the total observed molecular variability can be explained by the influence of *in vitro* culture conditions on micropropagated samples. Estimation of the molecular distance (PhiPT) between both types of propagules showed that the observed separation was significant for all genotypes. Although micropropagation is generally considered to induce low levels of somaclonal variation, our results are in concordance with previous studies in multiple species, e.g. cassava [6], grapevine [34], hop [35], tomato [36], triticale [37], and wild barley [38]. PCoA and PCA also revealed a higher level of variability within micropropagated samples compared to field maintained samples, which supports the somaclonal origin of the observed variability. The observed high level of variability within micropropagated sweet potato ramets, would suggest that a high proportion of the detected variability is random in nature. According to Smulders and De Klerk [39], the reason for these random changes might be attributable to the extreme conditions exposed to tissue culture plants such as abnormal photoperiods, wounding, application of growth regulators, among others. These may lead to oxidative stress, which can cause epigenetic or genetic changes to the genome, leading to somaclonal variants [40]. Interestingly, our results also show that the variability acquired by micropropagated ramets from all three genotypes occupied similar Eigen space in relation to their field maintained counterparts. This indicates that the observed somaclonal variability is not entirely random as previously seen in other species e.g. cocoa [7, 8].

### Somaclonal variation is mainly driven by DNA methylation polymorphisms

To determine the nature (genetic or epigenetic) of the detected somaclonal variation, we first identified the number of methylation sensitive and of non-methylated loci (MSL and NML respectively) using the *msap* v.3.3.1 R package. Between 62 and 70% (depending on the genotype/primer combination analysed) of all analysed loci were MSL. This level of methylation sits within the range of those previously described for plant species [41] for the CG and CNG contexts present within the recognition site of *Hpa*II and *Msp*I. PCA analysis of MSL and NML showed separation between field-maintained and micropropagated plants, suggesting that both types of variation could be contributing to the *in vitro* culture induced differences in MSAP profiles detected. Pairwise analysis of the molecular distance (PhiST) between field maintained and micropropagated plants, showed higher distances for all three genotypes when MSL where used. Moreover, while differences in MSL frequencies between the propagation methods were statistically significant, they were not when calculated using NML. Taken collectively, this indicates that changes in DNA methylation, rather than genetic, accounted for most, if not all, of the variability observed.

We then compared the band patterns of samples restricted with *Hpa*II and *Msp*I in order to assess the directionally of DNA methylation change induced by *in vitro* culture (i.e. Hypermethylation vs hypomethylation). Micropropagated plants presented higher levels of fully methylated sites and of internal cytosine methylation and lower levels of unmethylated and hemimethylated sites. This suggests that *in vitro* culture induces a global increase in DNA methylation. Previous studies have also reported higher levels of DNA methylation in tissue culture regenerants, e.g. hypermethylation on banana *in vitro* propagated clones relative to conventional ones [42], higher ratios of fully methylated CCGG sites in grapevine somaclones [43], and increased global levels DNA methylation of meristem cultures of *Malus xiaojinensis* [44].

### Micropropagation induced somaclonal variation in sweet potato is genotype dependent

As discussed above, PCoA and PCA of MSAP profiles revealed that *in vitro* induced molecular variability shifted micropropagated samples from all three genotypes into the same Eigen space. This indicates that a significant portion of the *in vitro* culture-induced epialleles are shared by all plants independently of their genotype. However, our study also revealed that while Bohye and Otoo plants showed the most extensive epigenetic and genetic variability respectively, micropropagated plants from genotype Ogyefo, consistently showed to be the least affected by micropropagation, i.e. they presented 1. the lowest percentage of total variability explained by micropropagation; 2. the lowest levels of total somaclonal variation (PhiPT) and the lowest levels of both genetic and epigenetic variability (PhiST). Previous studies have shown that factors affecting the level and type of somaclonal variability include: micropropagated species, the ortet’s genetic background and GC content, culture type and duration, plant hormones used, tissue used as explant material, among others [45, 46]. Here, all genotypes were exposed to the same in vitro culture conditions for the same period of time, which indicates that the extent of molecular variability inflicted by sweet potato micropropagation is genotype dependent.

## Conclusions

Here we show that micropropagation reduces the nutritional value of sweet potato tubers and that micropropagated plants are both genetically and epigenetically dissimilar from field maintained plants. The higher levels of variability in the nutritional composition and of molecular diversity observed within micropropagated plants makes tempting the speculation that there is a direct relation between both. Regardless, the anonymous nature of the MSAP markers, used here to characterize somaclonal variation at a molecular level, does not allow us to asseverate that the detected DNA methylation polymorphisms are the drivers of the observed loss in nutritional value. Still, our results provide a useful start point from which to assemble a more comprehensive picture of the functional role of *in vitro* culture induced DNA methylation changes affecting the nutritional value of biofortified crops. More importantly, since future sweet potato biofortification plans includes the use of *in vitro* culture to generate disease free propagules, our findings highlight the importance of including an assessment of the impact of micropropagation on nutritional values, with a special focus on beta-carotene content, of any novel biofortified sweet potato cultivar prior to their commercial release, to avoid the catastrophic costs to the industry previously seen with other *in vitro* propagated crops [47].

## Acknowledgements

Thanks to the staff of the sweet potato breeding programme, molecular biology, and tissue culture at CSIR-Crops Research Institute. The contribution of the staff of Environmental Epigenetics & Genetics Group as well as ARC Centre of Excellence Plant Energy Biology, of the School of Agriculture, Food and Wine, University of Adelaide, is much appreciated. The Bill & Melinda Gates Foundation through the International Centre for Genetic Engineering and Biotechnology (ICGEB), as a capacity building initiative in biotechnology regulation in sub-Saharan Africa, funded this research. The findings and conclusions contained in this publication are those of the authors and do not necessarily reflect positions or policies of the Bill & Melinda Gates Foundation nor the ICGEB. ICGEB participated in the project initial design but had not involvement during the data collection, analysis and manuscript drafting. The submission and publication of this manuscript was approved by ICGEB.

## Supporting Information

**Fig S1. Experimental layout**. Micropropagated (shown in orange border) and field-maintained plants (black bordered) planted in a common garden in a Randomized Complete Block Design with three replicates. Each block consisted of three plots: Bo (Bohye), Og (Ogyefo), and Ot (Otoo). Two Ridges (4.5 m long), spaced at 1 m, were made on each plot and fourteen vine cuttings of about 30 cm were planted on each ridge, with an interval of 30 cm between plants. The blocks were bordered with guard rows to reduce the effects of biotic and abiotic factors on the edge rows.

**Fig S2. PCoA results of the 12 primer combinations used for MSAP pilot studies**.

**Fig S3. Analysis of molecular somaclonal variation induced by micropropagation of sweet potato. A)** PCoA generated using GenAlex v6.5 from MSAP profiles from micropropagated (empty symbols) and field-maintained (full symbols) plants from genotypes of Bohye (green), Ogyefo (red), and Otoo (blue) (n = 24). MSAP profiles were amplified from genomic DNA restricted using *Hpa*II (circles) or MspI (squares) and amplified using primer combinations E and I (Results were calculated using loci from both primer combinations together). **B)** PCA generated by msapR 3.3.1 using MSAP profiles as above. Label on centroids indicate genotype (i.e. Bohye (Bo), Ogyefo (Og), and Otoo (Ot)) and type of propagule (i.e. Field maintained (FM) and micropropagated (VF)).

**Fig S4. Analysis of epigenetic and genetic variability induced by micropropagation in three sweet potato genotypes**. PCA generated by *msap*R 3.3.1 from MSAP profiles from micropropagated (blue symbols) and field-maintained (red symbols) plants from genotypes of Bohye (a and d), Ogyefo (b and e), and Otoo (c and f) (n=24). MSAP profiles were amplified from genomic DNA restricted using *Hpa*II and *Msp*I (profiles combined for analysis) and amplified using primer combinations E (a-c) and I (d-f). Epigenetic variability was calculated using Methylation Sensitive Loci (a1-f1) and genetic variability using Non-Methylated Loci (a2-f2). Plants were grown on a Randomized Complete Block Design with three replicated blocks and 24 plants/block/genotype/propagation system.

**Table S1. Identity, origin, year of release, and preferred ecology of the three sweet potato genotypes (Bohye, Ogyefo, and Otoo) used in the study**. block/genotype/propagation system.

**Table S2** Scale of reference (1-9) and definition of scores for virus incidence, foliar and root morphological descriptors.

**Table S3**. Oligonucleotides used during Methylation Sensitive Amplified Polymorphisms protocol with their sequences and function. Primer selective bases are highlighted in bold.

**Table S4**. Results of **s**elective primer combinations for MSAP pilot study. Primers selected are indicated with an asterisk. The number of alleles (# of loci), percentage reproducibility of alleles (% Rep), and number of differential alleles (# diff. alleles) are displayed.

**Table S5**. Mean foliage and root quality phenotypic scores for micropropagated (M) and field-maintained (F) populations of three sweet potato genotypes Bohye, Ogyefo, and Otoo. ILC=Immature Leaf Colour, MLC=Mature Leaf Colour, ALVP=Abaxial Leaf Vein Pigmentation, PVC=Predominant Vine Colour, SVC=Secondary Vine Colour, PP=Petiole Pigmentation, PT=Plant Type, M=micropropagated, and F=Field-maintained plants.

